# Expression of Tacr1 and Gpr83 by spinal projection neurons

**DOI:** 10.1101/2025.03.19.643956

**Authors:** Wenhui Ma, Allen C. Dickie, Erika Polgár, Mansi Yadav, Raphaëlle Quillet, Maria Gutierrez-Mecinas, Andrew M. Bell, Andrew J Todd

**Author notes:** These authors contributed equally (Allen C. Dickie, Erika Polgár). Email addresses: WM, ACD, EP, MY, RQ, MG-M, AMB, AJT.

## Abstract

Anterolateral system (ALS) projection neurons underlie perception of pain, itch and skin temperature. These cells are heterogeneous, and there have therefore been many attempts to define functional populations. A recent study identified two classes of ALS neuron in mouse superficial dorsal horn (SDH) based on expression of the G protein-coupled receptors Tacr1 or Gpr83. It was reported that cells expressing these receptors formed largely non-overlapping populations, and that ∼60% of ALS cells in SDH expressed Tacr1. An additional finding was that while Tacr1- and Gpr83-expressing ALS cells projected to several brain nuclei, their axons did not reach the ventral posterolateral (VPL) thalamic nucleus, which is reciprocally connected to the primary somatosensory cortex. These results were surprising, because we had reported that ∼90% of SDH ALS neurons in the mouse possess the neurokinin 1 receptor (NK1r), which is encoded by Tacr1, and in addition the VPL is thought to receive input from lamina I ALS cells. Here we use retrograde and anterograde labelling in Tacr1^CreERT2^ and Gpr83^CreERT2^ mice to reinvestigate the expression of the receptors by ALS neurons and to reassess their projection patterns. We find that ∼90% of ALS neurons in SDH express Tacr1, with 40-50% expressing Gpr83. We also show that axons of both Tacr1- and Gpr83-expressing ALS neurons reach the VPL. These results suggest that ALS neurons in the SDH that express these GPCRs show considerable overlap, and that they are likely to contribute to the sensory-discriminative dimension of pain through their projections to VPL.

## Introduction

Projection neurons belonging to the anterolateral system (ALS) transmit somatosensory information from the spinal cord to the brain, and are required for perception of pain, itch and skin temperature [1–5]. These cells are located in various regions of the spinal cord, including the superficial dorsal horn (SDH; laminae I and IIo), the lateral spinal nucleus (LSN) and the deeper laminae (in particular, lamina V). They project to several brain regions, including the thalamus, periaqueductal grey matter (PAG), lateral parabrachial area (LPB) and certain other brainstem nuclei [1,4]. Physiological studies have shown that ALS projection neurons are heterogeneous, and there have therefore been several attempts to define functional populations and identify molecular markers that would allow these to be targeted [6–8].

All spinal projection neurons in lamina I-IIo belong to the ALS [1,9], and they have attracted particular interest because many of these cells are densely innervated by nociceptive primary afferents, and the majority respond to noxious stimuli [10–15]. A recent study by Choi et al [6] identified two genetic markers for lamina I-IIo ALS neurons. One of these was Tacr1, the gene that encodes the neurokinin 1 receptor (NK1r), and this choice was based on previous studies showing that the receptor is expressed by the majority of lamina I projection neurons in the rat [10,16,17] and mouse [18]. The other marker, identified from a screen of BAC transgenic lines, was Gpr83, which codes for an orphan receptor [6]. Choi et al focused on ALS neurons in laminae I-IIo, and using newly developed mouse lines (Tacr1^CreERT2^ and Gpr83^CreERT2^) reported that these receptors were expressed in largely non-overlapping populations that accounted for ∼88% of spinoparabrachial neurons in the superficial laminae, with ∼57% expressing Tacr1 and ∼45% expressing Gpr83 [6].

However, we had previously found that the proportion of lamina I spinoparabrachial neurons that expressed the NK1r in the mouse was considerably higher (∼90%) [18], raising the possibility that some neurons with the receptor were not captured in the Tacr1^CreERT2^ mouse, and therefore that the extent of overlap of Tacr1 and Gpr83 had been underestimated. Furthermore, our recent transcriptomic analysis of Phox2a-derived ALS neurons identified 5 distinct clusters (ALS1-5) among these cells, but did not find that either Tacr1 or Gpr83 was specifically associated with any of these clusters [8]. Another unexpected finding of the study of Choi et al [6] was that although axons of both Tacr1- and Gpr83-expressing projection neurons could be followed to medial and posterior nuclei of the thalamus, they did not appear to innervate the ventral posterolateral (VPL) nucleus of the thalamus [6], even though VPL is thought to be a target for lamina I ALS neurons [4,19]. VPL is reciprocally connected with the primary somatosensory cortex, and ALS projections to this nucleus are therefore likely to contribute to the sensory-discriminative dimension of pain [3,4].

The aims of the present study were therefore to reassess the expression of Tacr1 and Gpr83 among spinal projection neurons, and to investigate the projection patterns of their axons. Specifically, we tested the hypotheses that Tacr1 and Gpr83 are expressed in overlapping populations in the SDH, and that the brain projections from Tacr1 and Gpr83 ALS cells include VPL.

## Materials and Methods

All animal experiments were approved by the Ethical Review Process Applications Panel of the University of Glasgow and were performed in accordance with the UK Animals (Scientific Procedures) Act 1986. The study was carried out in compliance with the ARRIVE guidelines.

### Animals and surgical procedures

We used the two mouse lines generated by Choi et al [6] in which Cre recombinase fused to the ligand-binding domain of the estrogen receptor was knocked into the Tacr1 (Tacr1^CreERT2^) or Gpr83 (Gpr83^CreERT2^) locus. For some experiments, these were cross-bred with the Rosa-CAG-LSL-tdTomato (Ai9) reporter line [20].

To determine the proportion of retrogradely labelled projection neurons in laminae I-IIo that were tdTomato-positive in the Tacr1^CreERT2^;Ai9 and Gpr83^CreERT2^;Ai9 crosses, we used tissue from 2 mice for each cross that had received injections of cholera toxin B subunit (CTB) into the caudal ventrolateral medulla (CVLM) as part of a previous study [21]. Tamoxifen had been administered to these mice as described by Choi et al (1.5 mg for Tacr1^CreERT2^;Ai9, 2.5 mg for Gpr83^CreERT2^;Ai9, given by intraperitoneal [i.p.] injection at between 21 and 30 days of age). In addition, two mice of each cross that had been treated with tamoxifen in the same way received an injection of CTB (300 nl, 0.2% or 0.5%) targeted on the LPB, as described previously [18]. These animals were anaesthetised with isoflurane (∼1.5%) and received perioperative analgesia (buprenorphine 0.1 mg/kg and carprofen 10 mg/kg, s.c.). The mice were placed in a stereotaxic frame and injections were made through burr holes in the skull, using glass micropipettes. These were left in place for 2 minutes after completion of the injection to avoid leakage of CTB up the track. The mice were re-anaesthetised 2 days later (20 mg pentobarbitone i.p.) and perfused through the left cardiac ventricle with fixative that contained 4% freshly depolymerised formaldehyde.

Anterograde tracing to reveal the axons of the projection neurons was performed by injecting AAV.Cre^ON^.tdTomato into the spinal cords of 4 Tacr1^CreERT2^ and 3 Gpr83^CreERT2^ mice (Table 1). In these cases, 3 mg of tamoxifen was administered by i.p. injection on two consecutive days, starting at least 4 days after the operation. The intraspinal injections were performed as described previously [21–24]. The mice were anaesthetised and received perioperative analgesia as described above. They were placed in a stereotaxic frame and after surgical exposure the T12 and L1 vertebrae were clamped. Injections were performed through glass micropipettes (tip diameter ∼30-40 μm) attached to a 10 μL Hamilton syringe on the right side of the spinal cord into either the L3 segment, or the L3, L4 and L5 segments. Injections into L3 and L5 were made through the intervertebral spaces on either side of the T13 vertebra, while those into L4 were made through a small hole drilled in the lamina of the T13 vertebra. Each injection consisted of 300 nl of AAV.Cre^ON^.tdTomato containing 9.48 × 10^7^ gene copies (AAV1 serotype, CAG promoter, VVF Zurich v167-1) and was administered at a rate of 30-40 nl/min. The pipette was left in place for 5 mins to minimise leakage of the injectate. The mice survived for between 35-43 days after tamoxifen injection and were then deeply anaesthetised and perfused with fixative as described above.

**Table 1.**
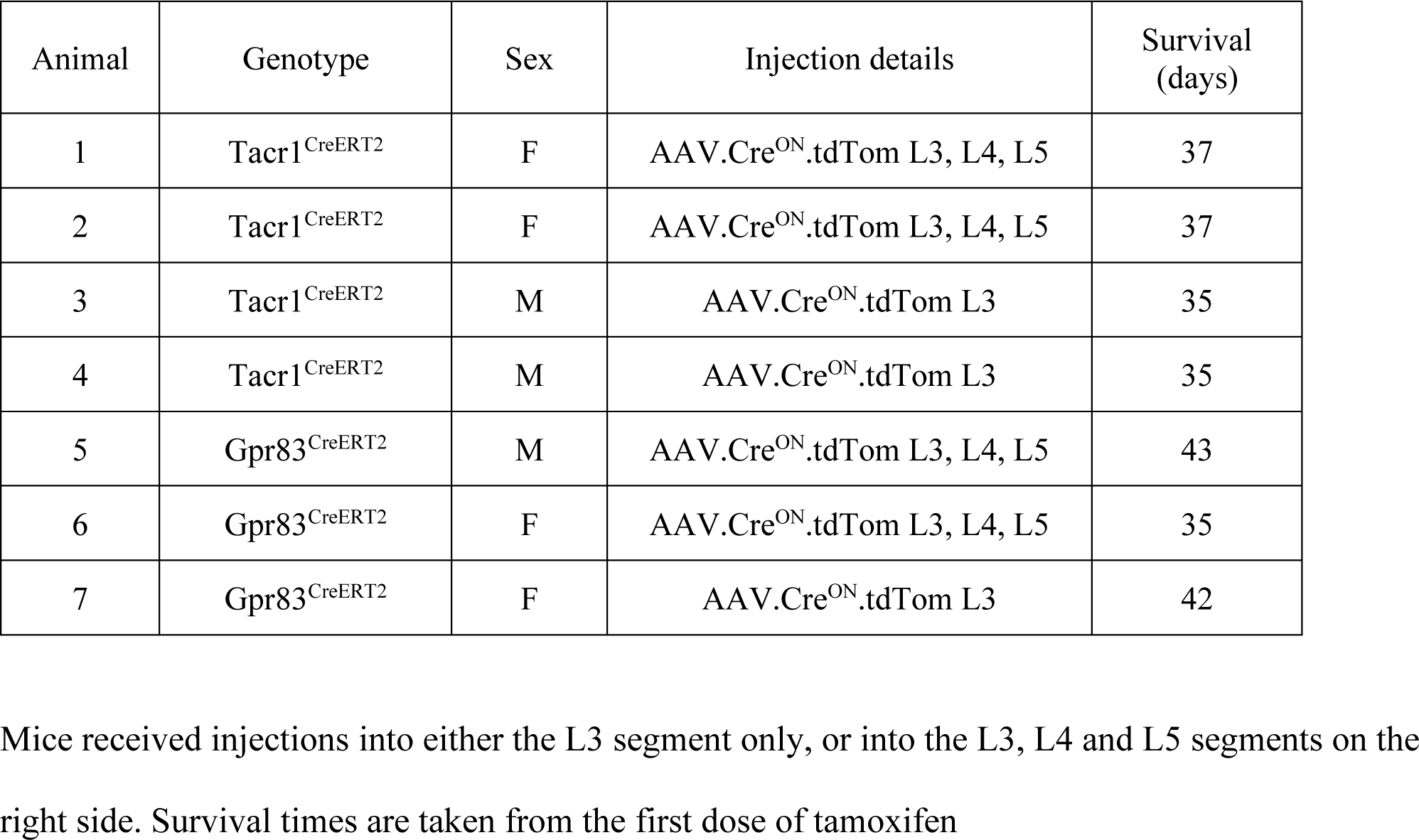
Experiment details for mice used in viral anterograde tracing experiments.

| Animal | Genotype | Sex | Injection details | Survival (days) |
| --- | --- | --- | --- | --- |
| 1 | Tacr1<sup>CreERT2</sup> | F | AAV.Cre<sup>ON</sup>.tdTom L3, L4, L5 | 37 |
| 2 | Tacr1<sup>CreERT2</sup> | F | AAV.Cre<sup>ON</sup>.tdTom L3, L4, L5 | 37 |
| 3 | Tacr1<sup>CreERT2</sup> | M | AAV.Cre<sup>ON</sup>.tdTom L3 | 35 |
| 4 | Tacr1<sup>CreERT2</sup> | M | AAV.Cre<sup>ON</sup>.tdTom L3 | 35 |
| 5 | Gpr83<sup>CreERT2</sup> | M | AAV.Cre<sup>ON</sup>.tdTom L3, L4, L5 | 43 |
| 6 | Gpr83<sup>CreERT2</sup> | F | AAV.Cre<sup>ON</sup>.tdTom L3, L4, L5 | 35 |
| 7 | Gpr83<sup>CreERT2</sup> | F | AAV.Cre<sup>ON</sup>.tdTom L3 | 42 |

Mice received injections into either the L3 segment only, or into the L3, L4 and L5 segments on the right side. Survival times are taken from the first dose of tamoxifen

### Immunohistochemistry, confocal scanning and analysis

Immunohistochemical reactions were performed as described previously [25] on 60 μm thick transverse sections of spinal cord cut with a vibrating blade microtome (Leica VT1200 or VT1000), or on 50 or 100 μm thick coronal sections of brain cut with the vibrating blade microtome or with a cryostat (Leica CM1950). The sources and concentrations of antibodies used are listed in Table 2. Sections were incubated for 3 days at 4°C in primary antibodies diluted in phosphate buffered saline that contained 0.3M NaCl, 0.3% Triton X-100 and 5% normal donkey serum, and then overnight in appropriate species-specific secondary antibodies that were raised in donkey and conjugated to Alexa 488, Alexa 647 or Rhodamine Red (Jackson Immunoresearch) or to Alexa 555-plus (Thermo Fisher Scientific). All secondary antibodies were used at 1:500 (in the same diluent), apart from those conjugated to Rhodamine Red, which were diluted to 1:100. Following the immunohistochemical reaction, sections were mounted in anti-fade medium and stored at −20°C. Transverse spinal cord sections through the L4 segments of the mice that had received injections of CTB into the brainstem were reacted with antibodies against mCherry, CTB and NK1r. Sections through the spinal cord and brain from the mice that received intraspinal injections of AAV.Cre^ON^.tdTomato were reacted with antibodies against mCherry and monoclonal NeuN antibody. Sections from the brains of mice that had received injections targeted on the LPB were reacted to reveal CTB using an immunoperoxidase method [18,21].

**Table 2.** Antibodies used in this study.

| Antigen | Species | Dilution | Source | Catalogue # | RRID |
| --- | --- | --- | --- | --- | --- |
| mCherry* | Rat | 1:1000 | Invitrogen | M11217 | RRID:AB_2536611 |
| CTB† | Goat | 1:5000 | List Biological | 703 | RRID:AB_10013220 |
| NK1r | Rabbit | 1:1000 | Sigma-Aldrich | S8305 | RRID:AB_261562 |
| NeuN-Alexa647 | Rabbit | 1:1000 | Abcam | ab190565 | RRID:AB_2732785 |
| NeuN | Guinea pig | 1:1000 | Synaptic Systems | 266 004 | RRID:AB_2619988 |

\*The mCherry antibody recognises tdTomato; †The CTB antibody was used at 1:100,000 for immunoperoxidase staining of brain injection sites

For analysis of retrograde labelling, sections of spinal cord were scanned with a Zeiss 710 LSM (Argon multi-line, 405 nm diode, 561 nm solid state and 633 nm HeNe lasers) through a 40× oil-immersion objective (numerical aperture 1.3) with the confocal aperture set to 1 Airy unit or less, to generate z-series through the full thickness of the section, and to include the entire width of the SDH. The resulting scans were analysed with Neurolucida for Confocal software (MBF Bioscience). Initially, the channel corresponding to CTB was viewed, and all CTB-labelled cells located in lamina I or IIo for which at least part of the nucleus (seen as a filling defect) was visible were plotted onto an outline of the dorsal horn. The other two channels were then switched on, and the presence or absence of tdTomato and/or NK1r immunoreactivity was recorded for each CTB-labelled cell.

For the anterograde tracing study sections were initially viewed on a fluorescence microscope (Zeiss Axioscope 5) and selected sections were scanned with the confocal microscope through 10× (NA 0.3) and 20× (NA 0.8) dry lenses, and the 40× oil-immersion lens. Identification of brain regions that contained tdTomato-labelled axons was based on the atlas of Franklin and Paxinos [26], except for the LPB, for which we used the same terminology as Choi et al [6], to allow for direct comparison with their study.

### Characterisation of antibodies

The sources and dilutions of primary antibodies used in the study are listed in Table 2. The mCherry antibody was raised against recombinant full-length protein, and the distribution of immunostaining matched that of native tdTomato. Specificity of the CTB antibody is demonstrated by the lack of staining in regions that did not contain injected or transported tracer. The NK1r antibody was raised against amino acids 393-407 of the rat NK1r and recognises a 46 kDa band in Western blots of rat brain extracts. It has been shown that there is no staining with this antibody in tissue from mice in which the NK1r has been deleted [27]. The NeuN antibodies give an identical staining pattern to that of a well-characterised mouse monoclonal NeuN antibody [28].

## Results

### Retrograde tracing experiments

CTB injection sites in the CVLM experiments were illustrated in Fig S3 of Ma et al [21], while LPB injection sites are shown in Fig S1.

The data obtained from the retrograde labelling studies are presented in Tables 3 and 4, and examples of immunostaining are shown in Figs 1 and 2. In both Tacr1^CreERT2^;Ai9 and Gpr83^CreERT2^;Ai9 mice, many tdTomato-positive cells were present throughout the upper part of the dorsal horn, as described previously [6]. In both CVLM and LPB experiments, CTB-labelled cells were particularly numerous in the SDH and present at a lower density in other regions of the spinal cord, with a pattern similar to that seen following equivalent tracer injections in the rat [29].

**Figure 1.**
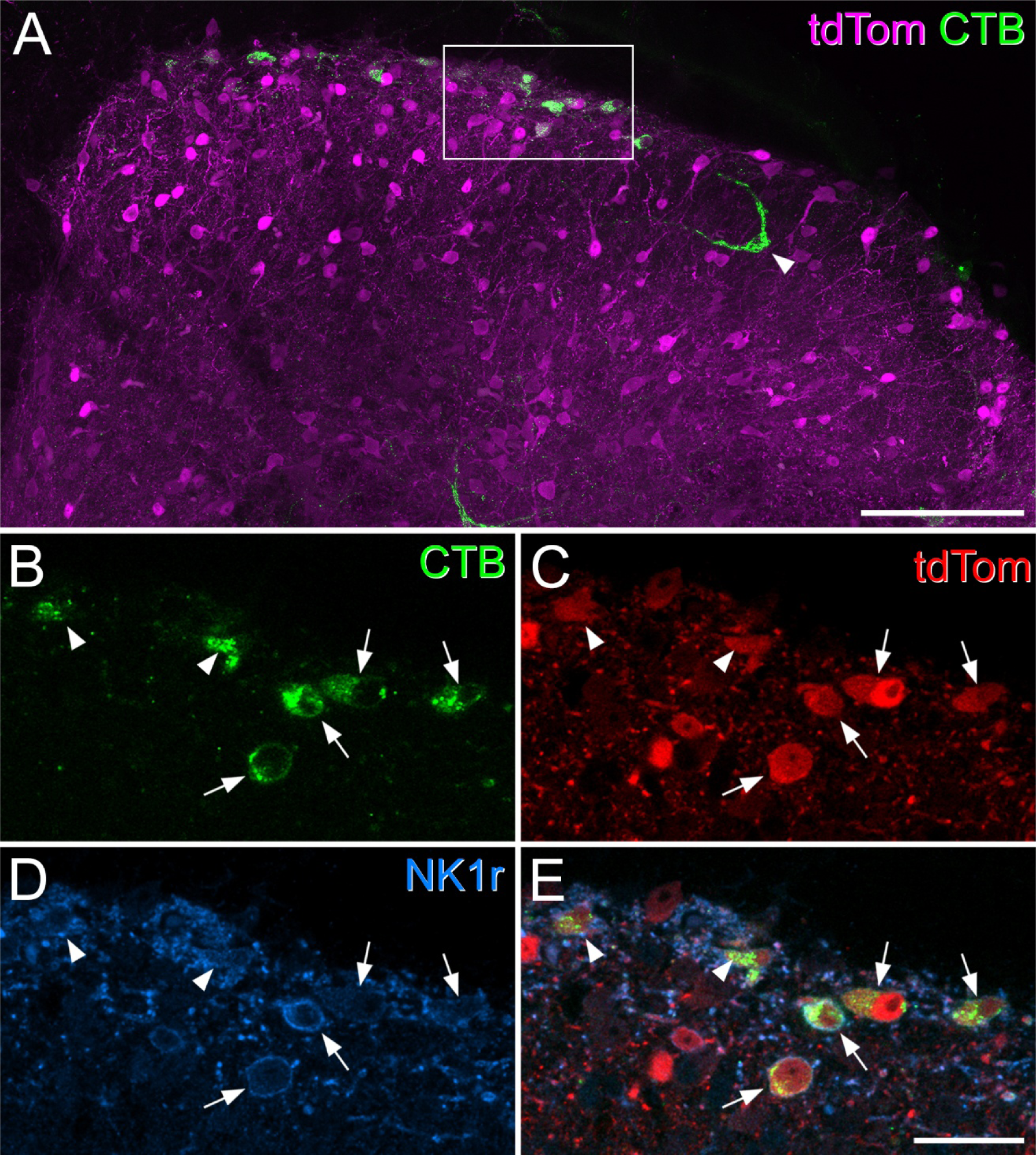
Most superficial dorsal horn projection neurons express the NK1 receptor and are tdTomato-positive in Tacr1^CreERT2^;Ai9 mice. ***A***, A transverse section through the upper part of the dorsal horn from the L4 segment of a Tacr1^CreERT2^;Ai9 mouse that had received an injection of CTB into the CVLM, scanned to reveal tdTomato (tdTom, magenta) and CTB (green). Several CTB-labelled cells are present in laminae I and IIo, and one is also seen in the inner part of lamina II (arrowhead). The box indicates the area shown at higher magnification in ***B***-***E***. ***B***-***E***, A scan (single optical section) through the region indicated in the box in ***A*** showing CTB (green), tdTomato (tdTom, red) and the NK1 receptor (NK1r, blue). Four CTB-labelled cells have their somata within this plane (arrows), while parts of two other cells (arrowheads) are also visible. All of these cells contain tdTomato and show NK1r-immunoreactivity in their membranes. ***A*** is a projection of 19 optical sections at 2 μm z-spacing. Scale bars: 100 μm (***A***) and 25 μm (***B***-***E***).

**Figure 2.**
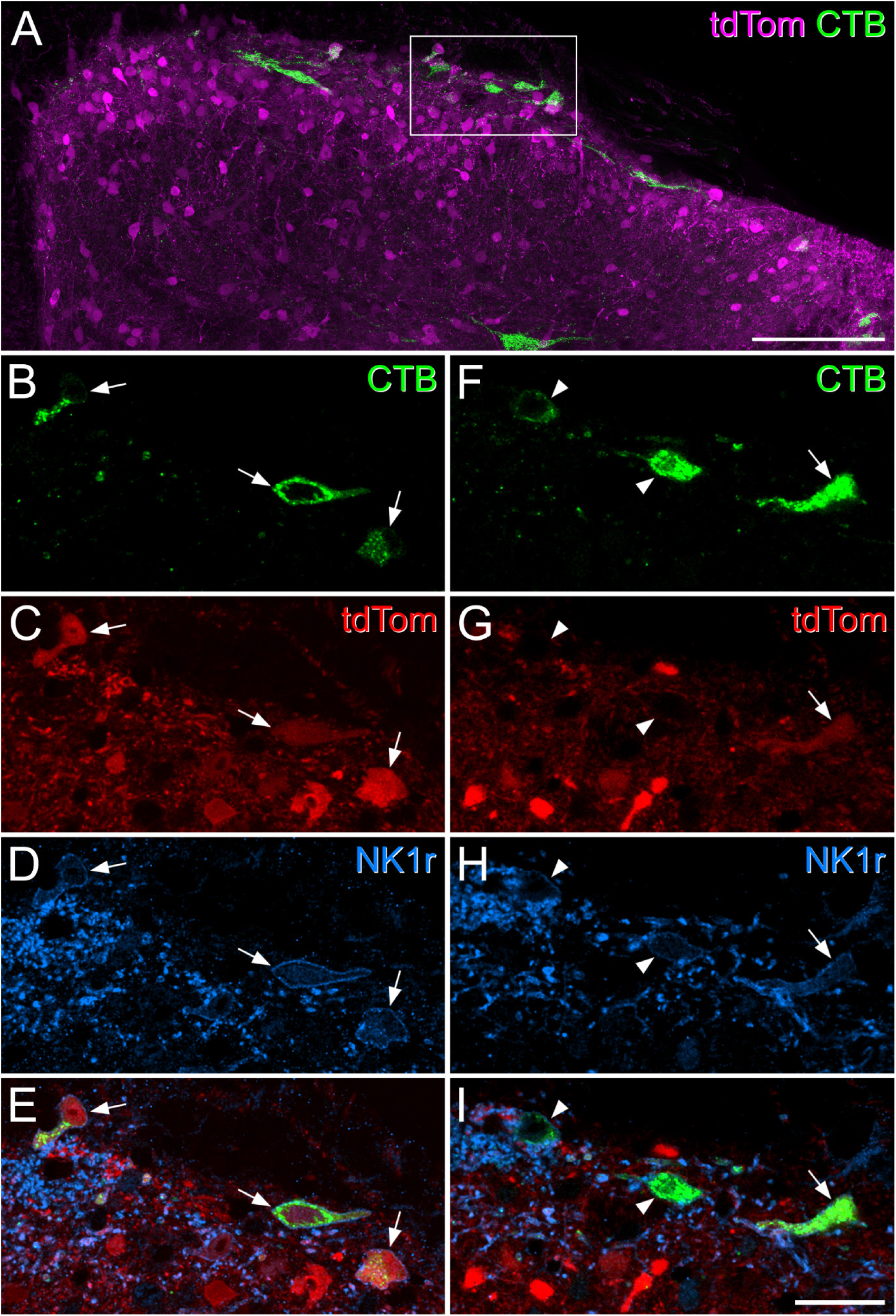
Expression of tdTomato and the neurokinin 1 receptor in superficial dorsal horn projection neurons in a Gpr83^CreERT2^;Ai9 mouse. ***A***, A transverse section through the upper part of the dorsal horn from the L4 segment of a Gpr83^CreERT2^;Ai9 mouse that had received an injection of CTB into the CVLM, scanned to reveal tdTomato (tdTom, magenta) and CTB (green). Several CTB-labelled cells are present in the superficial dorsal horn. The box indicates the region shown at higher magnification in ***B***-***I***. ***B***-***E*** show a single optical section through the superficial part of the tissue section, corresponding to the boxed region in ***A***. There are 3 CTB-labelled cells (arrows) all of which contain tdTomato and show NK1r-immunoreactivity in their membranes. ***F***-***H***, a projection of 2 adjacent optical sections taken from deeper within the tissue section in the same boxed region from ***A***. Again, 3 CTB-labelled cells are visible. One of these (arrow) contains tdTomato, while the other two (arrowheads) do not. All 3 cells have NK1r-immunoreactivity in the membrane. ***A*** is a projection of 24 optical sections at 2 μm z-spacing. Scale bars: 100 μm (***A***) and 25 μm (***B***-***I***).

**Table 3.** CTb and TdTomato expression following injection of CTb into the CVLM.

| Genotype | CTb cells | tdTom+ NK1r+ | tdTom+ NK1r- | tdTom- NK1r+ | tdTom- NK1r- | % with tdTom | % with NK1r | % NK1r+ with tdTom | % tdTom+ with NK1r |
| --- | --- | --- | --- | --- | --- | --- | --- | --- | --- |
| Tacr1<sup>CreERT2</sup>;Ai9 (n=2) | 51<br>(48, 54) | 41.5<br>(38, 45) | 5.5<br>(5, 6) | 2<br>(2, 2) | 2<br>(1, 3) | 92<br>(89.6, 94.4) | 85.2<br>(83.3, 87) | 95.4<br>(95.7, 95) | 88.3<br>(88.2, 88.4) |
| Gpr83<sup>CreERT2</sup>;Ai9 (n=2) | 51<br>(55, 47) | 24<br>(21, 27) | 2<br>(1, 3) | 19.5<br>(19, 20) | 5.5<br>(5, 6) | 50.7<br>(46.8, 54.5) | 85.3<br>(85.1, 85.4) | 55<br>(57.4, 52.5) | 92.7<br>(90, 95.4) |

Columns 2-6 show the mean numbers of retrogradely labelled lamina I cells of each type identified in transverse sections through the L4 segment (with individual values in brackets). Columns 7-8 show the proportions of retrogradely labelled lamina I cells with tdTomato or NK1r immunoreactivity, while column 9-10 show the proportion of retrogradely labelled NK1r-positive lamina I cells with tdTomato and *vice versa*.

**Table 4.** CTb and TdTomato expression following injection of CTb into the LPB.

| Genotype | CTb cells | tdTom+ NK1r+ | tdTom+ NK1r- | tdTom- NK1r+ | tdTom- NK1r- | % with tdTom | % with NK1r | % NK1r+ with tdTom | % tdTom+ with NK1r |
| --- | --- | --- | --- | --- | --- | --- | --- | --- | --- |
| Tacr1<sup>CreERT2</sup>;Ai9 (n=2) | 64.5<br>(66, 63) | 49<br>(45, 53) | 8.5<br>(11, 6) | 6<br>(9, 3) | 1<br>(1, 1) | 89.2<br>(84.8, 93.6) | 85.4<br>(81.8, 88.9) | 89<br>(83.3, 94.6) | 85.1<br>(80.4, 89.8) |
| Gpr83<sup>CreERT2</sup>;Ai9 (n=2) | 55.5<br>(54, 57) | 21<br>(23, 19) | 2<br>(0, 4) | 29<br>(29, 29) | 3.5<br>(2, 5) | 41.5<br>(42.6, 40.4) | 90.3<br>(96.3, 84.2) | 41.9<br>(44.2, 39.6) | 91.3<br>(100, 82.6) |

Columns 2-6 show the numbers of retrogradely labelled lamina I cells of each type identified in transverse sections through the L4 segment. Columns 7-8 show the proportions of retrogradely labelled lamina I cells with tdTomato or NK1r immunoreactivity, while column 9-10 show the proportion of retrogradely labelled NK1r-positive lamina I cells with tdTomato and *vice versa*.

In laminae I-IIo of the Tacr1^CreERT2^;Ai9 mice we found that most cells that were NK1r-immunoreactive contained tdTomato, and that many of the tdTomato cells had NK1r immunoreactivity in the plasma membrane (Fig 1). However, some cells that were NK1r-immunoreactive lacked tdTomato, while some tdTomato-positive cells did not have detectable NK1r (Fig S2). Among the CTB-labelled cells in laminae I-IIo, between 89-95% of those identified as NK1r-immunoreactive contained tdTomato, while 85-88% of those with tdTomato were defined as NK1r-immunoreactive (Tables 3, 4). Analysis of retrograde labelling in these mice revealed that the great majority of CTB-labelled cells in laminae I-IIo (92% for CVLM injections, 89% for LPB injections) contained tdTomato, and that 85% of the retrogradely labelled cells were NK1r-immunoreactive (Fig 1, Tables 3, 4).

In Gpr83^CreERT2^;Ai9 mice, 51% of laminae I-IIo cells labelled from the CVLM, and 42% of those labelled from the LPB, contained tdTomato (Tables 3, 4, Fig 2). Again, a very high proportion (85-90%) of the retrogradely labelled cells were NK1r-immunoreactive. Consistent with this, we found that most (91-93%) of the retrogradely labelled cells that contained tdTomato were NK1r-immunoreactive (Tables 3, 4).

Although we did not analyse tdTomato expression of retrogradely labelled cells in other areas of the spinal cord in detail, we noted that the majority of CTB-labelled cells in the LSN were tdTomato-positive in Tacr1^CreERT2^;Ai9 mice, while around one-third of CTB-positive LSN cells contained tdTomato in the Gpr83^CreERT2^;Ai9 mice (Fig S3). In the medial part of laminae VI-VII we found that some CTB-labelled cells were tdTomato-positive in each cross, while very few of the retrogradely labelled cells in the medial part of lamina V contained tdTomato in either cross (Fig S3).

### Anterograde tracing

The pattern of labelling in spinal cord and brain following injection of AAV.Cre^ON^.tdTomato into the lumbar spinal cord was generally similar in the two genotypes. Labelling was consistent between mice of the same genotype, but as expected it was denser in mice in which injections were made into three spinal segments compared to those in which injections were made into only one segment.

In both genotypes there was extensive expression of tdTomato around the injection sites, which was largely limited to the dorsal horn on the injected side, and this became more restricted at rostral levels (Fig 3). In the cervical cord, many tdTomato-containing axons were present in the white matter on both sides. These were particularly numerous in the dorsal part of the lateral funiculus on each side. In addition, there were labelled axons in the ventral part of the lateral funiculus and the ventral funiculus, and these were much more common on the side contralateral to the spinal injection. In both genotypes, there were also many fibres within the gracile tract in the ipsilateral dorsal funiculus. Axon collaterals were seen in the grey matter at all levels examined, especially in a region extending from the lateral part of lamina V towards the central canal. These collaterals were much more numerous on the side ipsilateral to the spinal injection (Figs 3, S4).

**Figure 3.**
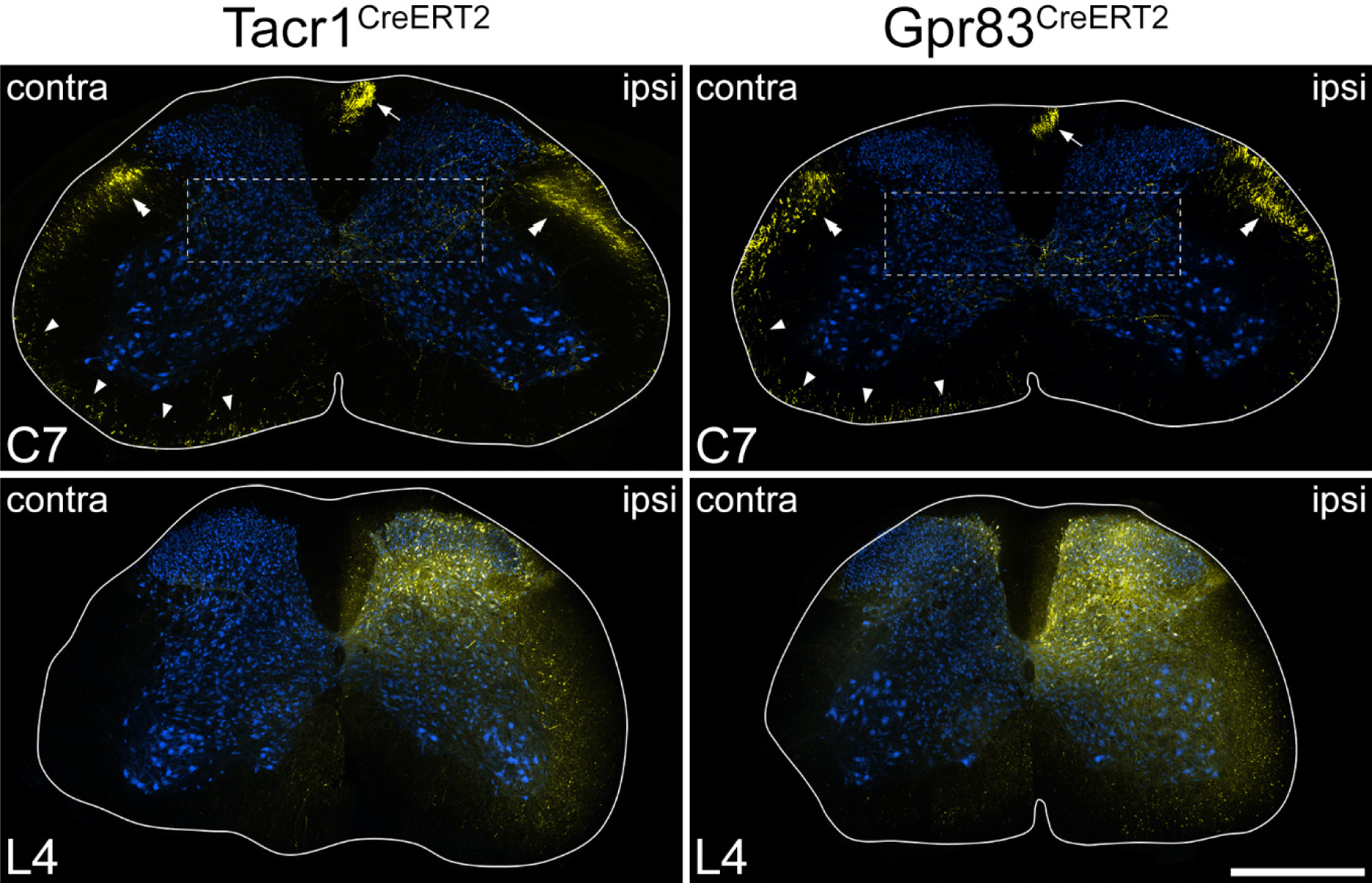
tdTomato labelling in the spinal cord in the anterograde tracing experiments. Transverse sections through the L4 and C7 segments of Tacr1^CreERT2^ and Gpr83^CreERT2^ mice that had received injections of AAV.Cre^ON^.tdTomato into the L3, L4 and L5 segments on one side. Sections have been immunoreacted to reveal tdTomato (yellow) and NeuN (blue). The white lines indicate the surface of the spinal cord. At the level of the injection site (L4) there is extensive labelling that is largely restricted to one side, although there is a very minor extension to the medial edge of the dorsal horn on the contralateral side in the Gpr83^CreERT2^ mouse. At the C7 level, there are numerous axons in various parts of the white matter. There is labelling in the gracile tract (arrows) and dense bundles of axons in the dorsal part of the lateral funiculus on each side (double arrowheads). In addition, there are labelled axons in the ventral part of the lateral funiculus, and in the ventral funiculus, particularly on the contralateral (contra) side (arrowheads). There are also collateral branches that innervate the deep grey matter on the ipsilateral (ipsi) side. Boxes indicate the areas shown at higher magnification in Fig S4 to reveal these collaterals. Scale bar = 500 μm.

In the medulla, the ascending bundles of labelled axons were located bilaterally near the ventral aspect (Fig 4), and collaterals extended from here into various reticular nuclei, as well as to the nucleus of the solitary tract (NTS). In both genotypes there was also dense terminal labelling in the medial nucleus of the inferior olive (IOM) on the contralateral side and in the ipsilateral gracile nucleus. The main bundle of ascending axons continued in a similar location near the ventral aspect of the brainstem until it reached a level corresponding to the rostral part of the pons, where it turned dorsally to reach the LPB. For both genotypes, axons passed around the lateral aspect of the LPB, giving collateral branches with boutons to the internal lateral (il), dorsal lateral (dl) and external lateral (el) nuclei (Figs 5 and S5). In addition, in the Tacr1^CreERT2^ mice there were projections to the central lateral (cl) nucleus, and a small projection to the ventral lateral (vl) nucleus, but fewer axons were seen in these nuclei in the Gpr83^CreERT2^ mice. The central lateral nucleus of the LPB (PBcl) was one of the few brain regions in which we observed a difference between the axonal projections of the Tacr1^CreERT2^ and Gpr83^CreERT2^ mice, and this finding was consistent with that of Choi et al [6] (their Fig 2). We have obtained further information concerning inputs to PBcl as part of a separate study in which we are investigating the projections of ALS cells that express the calcium-binding protein calbindin, using Calb1^Cre^ mice, and have found that these show strong labelling in the PBcl (Fig S5).

**Figure 4.**
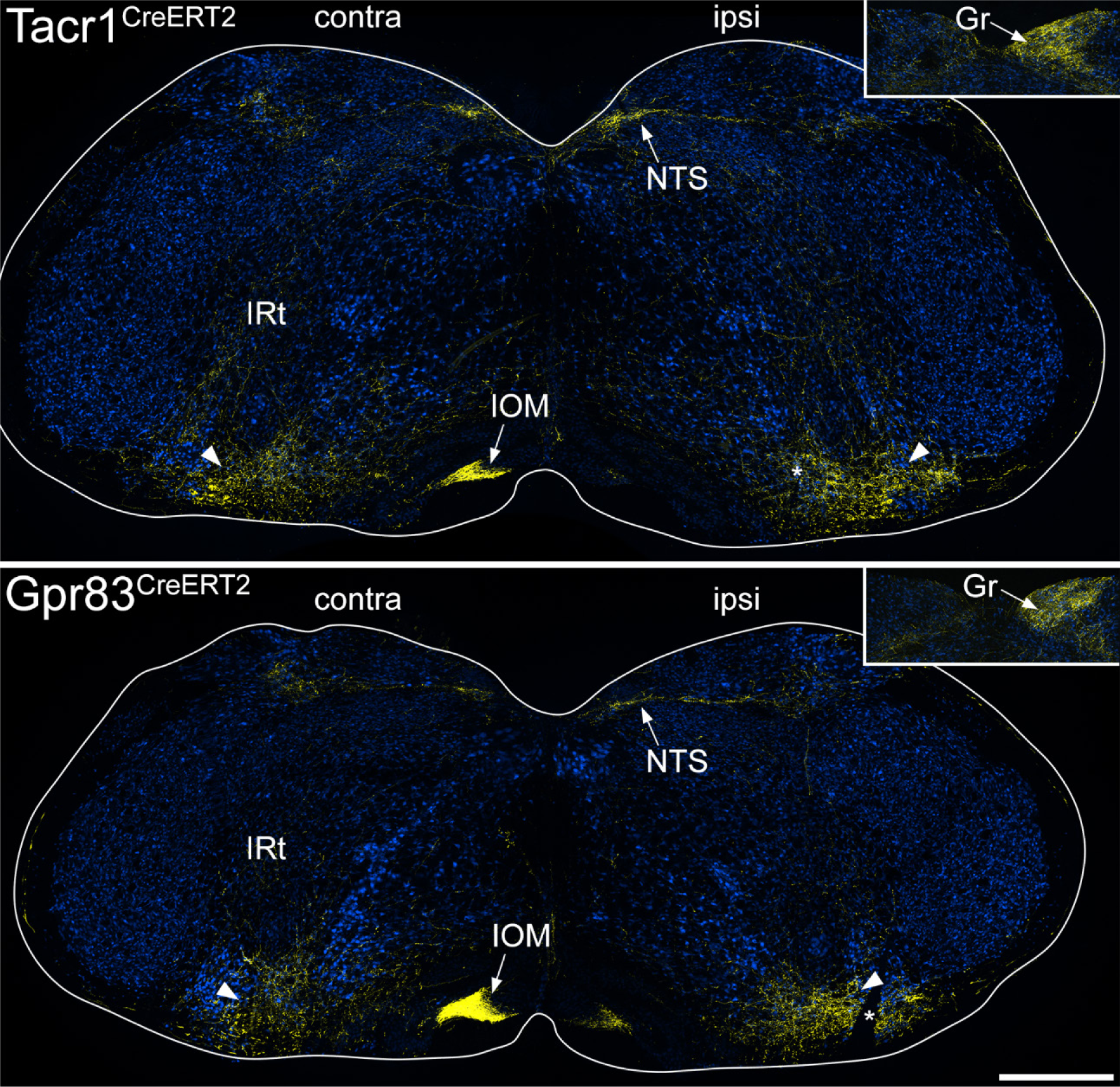
tdTomato labelling in the medulla in the anterograde tracing experiments. The main images show sections through the medulla at a level corresponding to ∼3.4 mm behind the interaural line from Tacr1^CreERT2^ and Gpr83^CreERT2^ mice that had received injections of AAV.Cre^ON^.tdTomato into the L3, L4 and L5 segments on one side. Sections have been immunoreacted to reveal tdTomato (yellow) and NeuN (blue). The sides contralateral (contra) and ipsilateral (ipsi) to the spinal injection are labelled on each image and the white lines indicate the surface of the medulla. Arrowheads show the position of the main ascending bundle on each side. From this, collateral branches innervate various regions, including the intermediate reticular nucleus (IRt) and the nucleus of the solitary tract (NTS) on both sides, and the medial nucleus of the inferior olive (IOM) on the contralateral side. The inset shows part of a section at a more caudal level through the medulla (∼ 4 mm caudal to the interaural line), with axons innervating the gracile nucleus (Gr) on the ipsilateral side. In each of the main images, the asterisk shows a cut made on the ipsilateral side that was used to orientate the sections. Scale bar = 500 μm.

**Figure 5.**
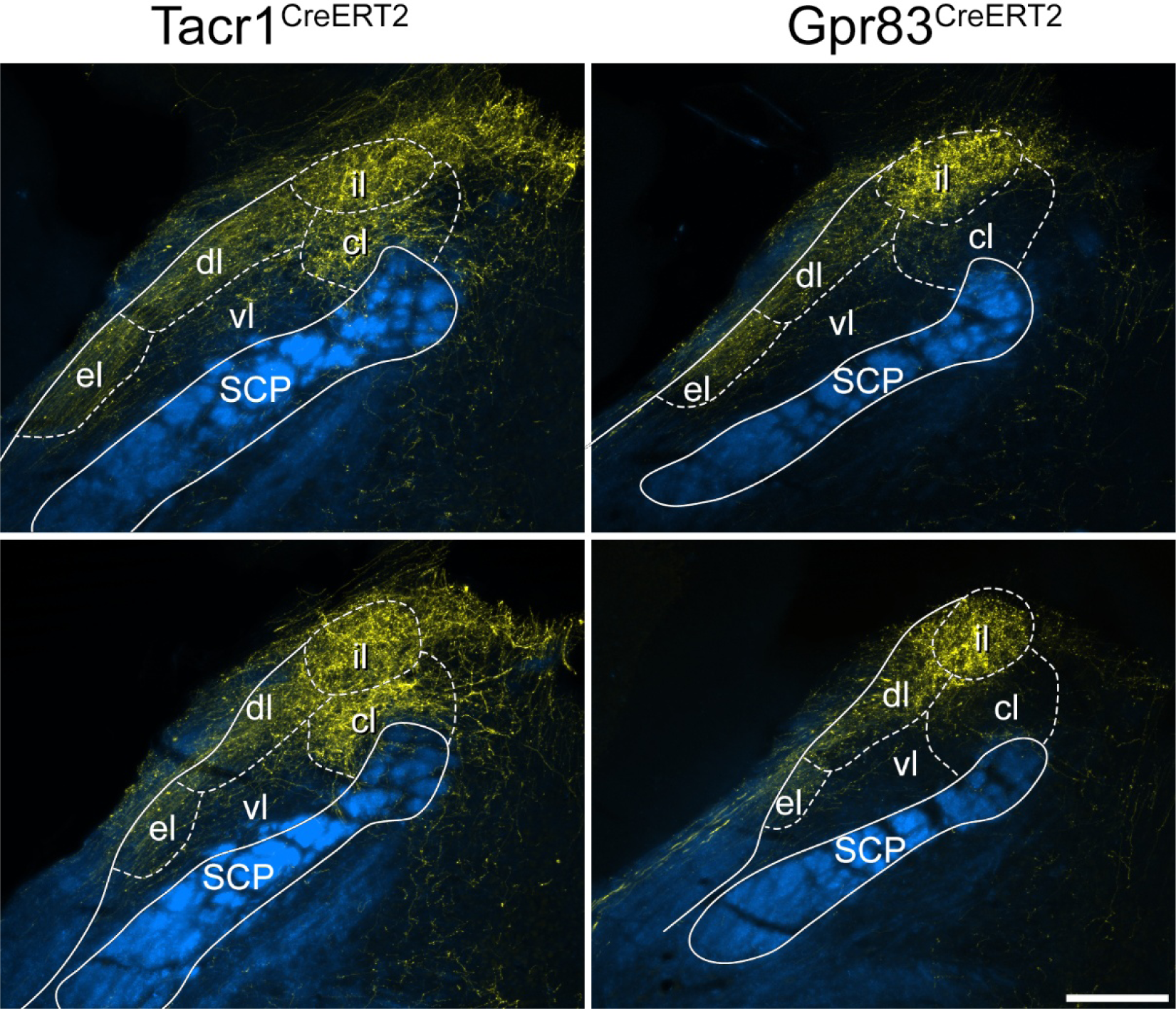
tdTomato labelling in the lateral parabrachial area in the anterograde tracing experiments. Images show sections through the contralateral lateral parabrachial (LPB) area from a Tacr1^CreERT2^ mouse and a Gpr83^CreERT2^ mouse that had received injections of AAV.Cre^ON^.tdTomato into the L3, L4 and L5 segments on one side. Immunolabelling for tdTomato (yellow) is superimposed on a dark-field image (blue). In each case, two serial sections through the brainstem are shown, with the lower image being from a slightly more caudal level. The superior cerebellar peduncle (SCP) is outlined for each section, and approximate positions of LPB nuclei are indicated: cl, central lateral; dl, dorsolateral; el, external lateral; il, internal lateral; vl ventral lateral. There is labelling in el, dl and il in both genotypes, as well as labelling in cl in the Tacr1^CreERT2^ mouse. Scale bar = 200 μm.

As described for ascending projections of somatostatin-expressing ALS neurons located in the lateral part of lamina V [21], some fibres crossed the midline in the cerebellar commissure (Fig S6). Some axons entered the cerebellar cortex, where they gave rise to large boutons that are presumably mossy fibre terminals, and these were more numerous for the Gpr83^CreERT2^ line. Further rostrally, axons arborised in the PAG, particularly the lateral part (LPAG), the posterior triangular (PoT) nucleus of the thalamus, and in both cases these were more numerous on the side contralateral to the injection (Fig 6). The most rostral projections were seen in the medial part of the thalamus, in particular the mediodorsal nucleus (MD), and in the contralateral ventral posterolateral (VPL) nucleus of the thalamus (Fig 7). Labelling in the contralateral VPL was similar in the two genotypes and occupied mainly the caudal part, extending from approximately 1.5 to 2.1 mm rostral to the interaural line.

**Figure 6.**
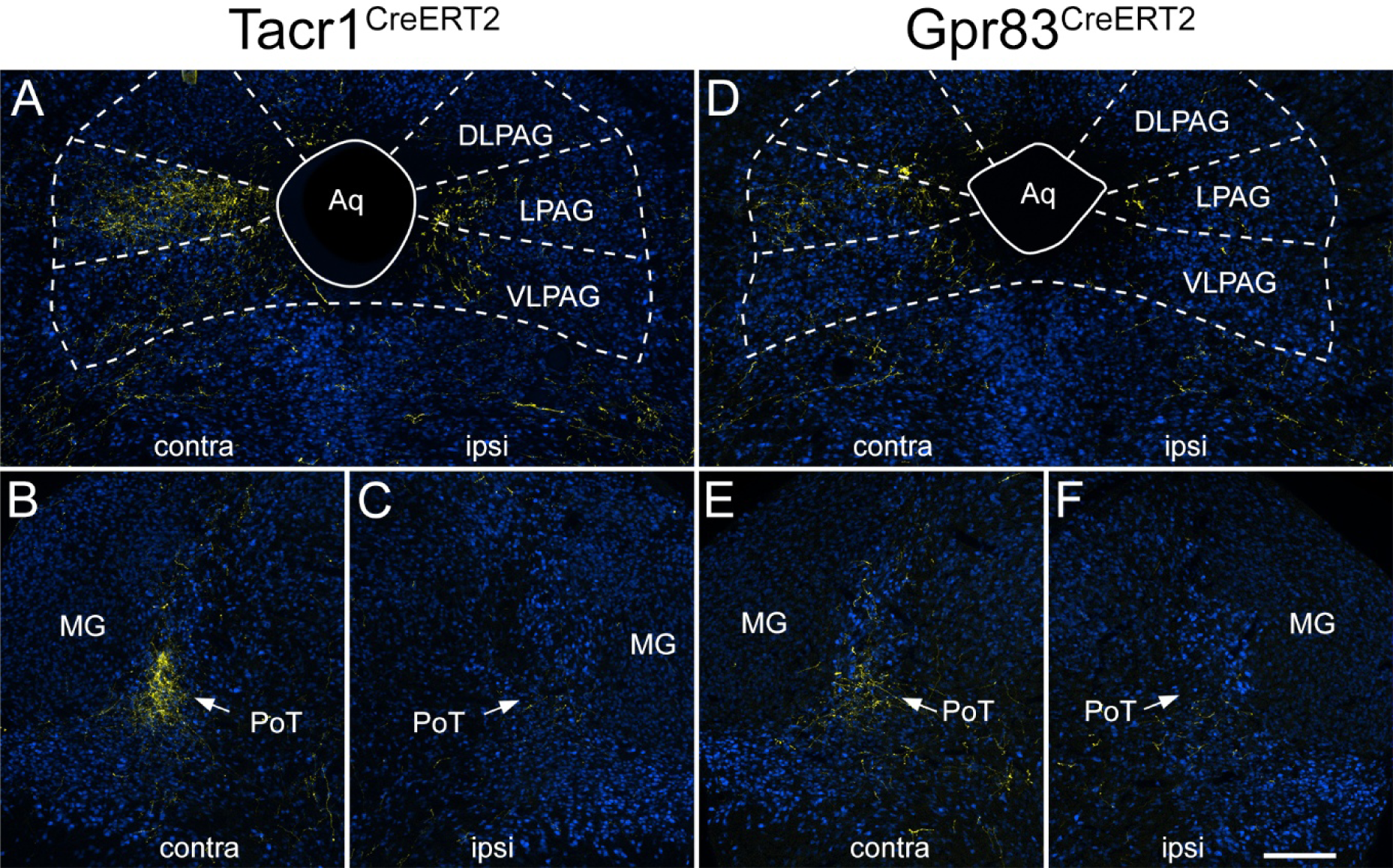
tdTomato labelling in the caudal thalamus and PAG in the anterograde tracing experiments. Images show sections from a Tacr1^CreERT2^ mouse (**A**-**C**) and a Gpr83^CreERT2^ mouse (**D**-**F**) that had received injections of AAV.Cre^ON^.tdTomato into the L3, L4 and L5 segments on one side. Sections have been immunoreacted to reveal tdTomato (yellow) and NeuN (blue). Sections correspond to levels approximately 0.8 mm behind (A, D) or 0.4 mm in front of (B, C, E, F) the interaural line. The sides contralateral (contra) and ipsilateral (ipsi) to the spinal injection are indicated on each image. **A**: tdTomato labelling in the PAG in the Tacr1^CreERT2^ mouse is denser on the contralateral side, and is mainly present in the lateral PAG (LPAG), with less labelling in dorsolateral (DLPAG) or ventrolateral (VLPAG) parts. There is some labelling in LPAG and VLPAG on the ipsilateral side. **B**, **C**: there is dense tdTomato-labelling in the posterior triangular (PoT) nucleus of the thalamus on the contralateral side in the Tacr1^CreERT2^ mouse, with minimal labelling on the ipsilateral side. The PoT can be recognised due to its location just medial to the medial geniculate nucleus (MG). **D**: PAG labelling in the Gpr83^CreERT2^ mouse is again denser on the contralateral side, and is mainly present in the LPAG. **E**, **F**: In the Gpr83^CreERT2^ mouse there is also labelling in the contralateral PoT. Scale bar = 200 μm.

**Figure 7.**
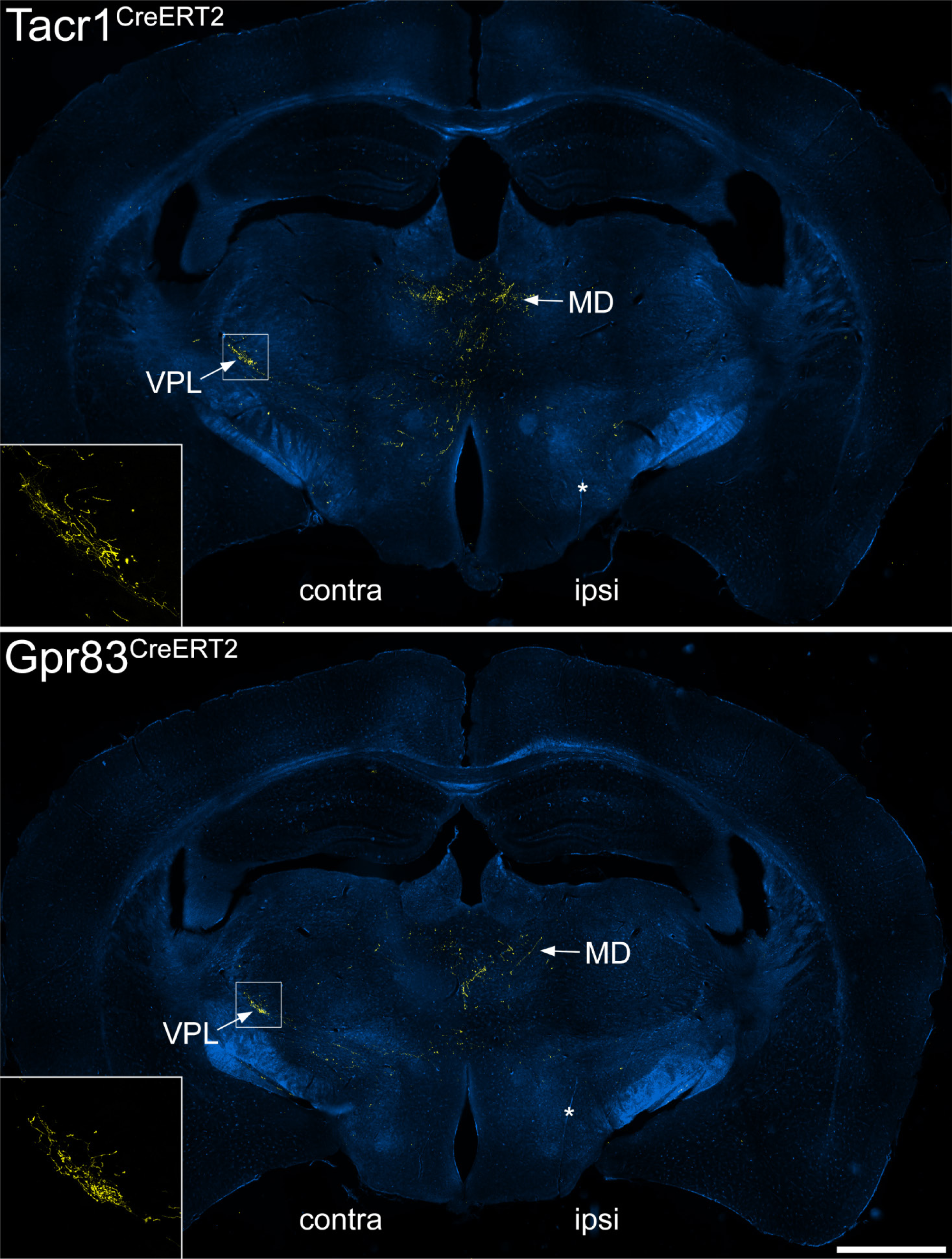
tdTomato labelling in the medial dorsal and ventral posterolateral thalamic nuclei in the anterograde tracing experiments. Immunolabelling for tdTomato (yellow) superimposed on a dark field image at a level ∼2.1 mm rostral to the interaural line from Tacr1^CreERT2^ and Gpr83^CreERT2^ mice that had received injections of AAV.Cre^ON^.tdTomato into the L3, L4 and L5 segments on one side. The sides contralateral (contra) and ipsilateral (ipsi) to the spinal injection are indicated on each image. Many tdTomato-labelled axons are present in the medial part of the thalamus, especially the mediodorsal nucleus (MD). In addition, in each case there is a cluster of axons in the ventral posterolateral nucleus (VPL) on the side contralateral to the injection site. The insets show the pattern of VPL labelling at higher magnification, and correspond to the boxes in the main parts of the figure. Asterisks shows cuts made on the ipsilateral side that were used to orientate the sections. Scale bar = 1 mm.

## Discussion

The main findings of this study are that: (1) the great majority (∼90%) of neurons in laminae I-IIo that are retrogradely labelled from the LPB or CVLM express tdTomato in the Tacr1^CreERT2^;Ai9 mouse; (2) most retrogradely labelled tdTomato-positive cells in laminae I-IIo of the Gpr83^CreERT2^;Ai9 mouse are NK1r-immunoreactive; (3) axonal projections of Cre-expressing cells in the Tacr1^CreERT2^ and Gpr83^CreERT2^ mice are broadly similar, and in both cases these include the VPL nucleus of the thalamus.

### Tacr1 and Gpr83 expression by spinal projection neurons

For the retrograde labelling analysis, we used tissue from mice that had received CTB injections into two different regions, CVLM and LPB. We had previously shown that when different tracers are injected into the corresponding sites in the rat, the great majority (>80%) of retrogradely labelled cells in lamina I contain both tracers [17]. It is therefore likely that a largely overlapping population of projection neurons in the SDH was captured from these two sites in the mouse.

The report that only ∼60% of lamina I-IIo spinoparabrachial neurons were Tacr1-positive [6] was very surprising, because we had previously found that ∼80% of lamina I projection neurons retrogradely labelled from various brain regions in rat were NK1r-immunoreactive [16,17,29–31], with an even higher proportion (90%) for lamina I spinoparabrachial neurons in the mouse [18]. Here we find that ∼90% of neurons in laminae I-IIo that were labelled from either CVLM or LPB express tdTomato in the Tacr1^CreERT2^;Ai9 mouse, consistent with our previous [18] and present immunohistochemical analyses. We also show that, at least for superficial projection neurons, there is a good correspondence between tdTomato and NK1r expression in this cross, suggesting that Cre expression in the Tacr1^CreERT2^ mouse reliably marks most cells that express the receptor. The most likely explanation for the difference between our findings and those of Choi et al [6] is therefore that recombination efficiency was lower in their experiments, and this probably reflects the well-established variability in tamoxifen-induced recombination when Cre^ERT2^ lines are crossed with reporter mice [24,32]. These results, together with the finding of tdTomato expression in projection neurons in other regions (LSN, deep dorsal horn, lateral lamina V) make it highly unlikely that Tacr1 expression defines a discrete population of ALS projection neurons.

In contrast, we found that 42-51% of retrogradely labelled lamina I-IIo neurons contained tdTomato in the experiments involving the Gpr83^CreERT2^;Ai9 mouse, and this is consistent the finding of Choi et al that Gpr83-expressing cells accounted for 45% of spinoparabrachial neurons in this region [6]. However, since the great majority of the tdTomato-containing projection neurons in our experiments were NK1r-immunoreactive, our findings do not support the suggestion that Tacr1 and Gpr83 are found in largely non-overlapping populations among the superficial projection neurons [6].

Our results also demonstrate that projection neurons captured in both genotypes are not restricted to the SDH. We had already shown that some ALS cells in lateral lamina V expressed Tacr1 or Gpr83 [21], and here we found tdTomato in some retrogradely labelled cells in other laminae, in particular the LSN. We also show that spinal injection of AAV.Cre^ON^.tdTomato in both genotypes resulted in axonal labelling in the ipsilateral gracile tract, with terminal labelling in the gracile nucleus. Tacr1 is not expressed in primary sensory neurons, and Gpr83 is apparently restricted to Trpm8-expressing afferents [33], which do not ascend in the dorsal columns. It is therefore likely that that gracile tract/nucleus labelling results from capture of neurons belonging to the postsynaptic dorsal column (PSDC) pathway in both genotypes.

### Axonal projections of Tacr1- and Gpr83-expressing projection cells

The distribution of anterogradely labelled axons within the cervical cord was complex, with labelling on both sides, and in various parts of the white matter. This probably results from the capture in both genotypes of several different projection neuron populations, including cells in the SDH, the LSN, lateral lamina V and the deep medial dorsal horn, as well as PSDC cells and a few spinocerebellar tract neurons. In addition, some of the axons seen in the cervical white matter probably belong to propriospinal neurons [21,34]. ALS cells project mainly to contralateral brain regions [4] and are therefore likely to have axons that ascend in the contralateral white matter. However, we recently reported that cells belonging to the ALS4 population [8], which are located in the lateral reticulated part of lamina V, had axons that ascended in the ipsilateral lateral funiculus, and gave extensive collateral branches to a region extending from lateral lamina V to the central canal at all spinal levels [21]. Since this population includes some cells with Tacr1 and/or Gpr83 [8,21], it is likely that some of the ipsilaterally-projecting axons and spinal cord collaterals seen in the cervical cord originate from cells in the lateral part of lamina V that belong to the ALS4 population. The relative sparsity of spinal collaterals on the contralateral side suggests that ALS cells with axons that ascend in the contralateral white matter give off fewer of these collaterals.

Within the brain, our findings show some similarities with those of Choi et al [6], since we saw projections to the IOM, the PAG, the PoT (which they refer to as MGm/SPFp) and medial thalamic nuclei in both genotypes. In agreement with Choi et al, we also found differences between the genotypes in projections to the LPB, with the PBcl nucleus receiving a dense input from axons expressing Tacr1, but not Gpr83. This suggests that a population of ALS neurons that are preferentially captured in the Tacr1^CreERT2^ mice project to the PBcl, and further insight into a possible source of these cells is provided by our finding that calbindin-expressing ALS neurons also project to PBcl. Calbindin is expressed in many ALS neurons in lamina I and the LSN [35], and these include lamina I cells belonging to our ALS2 and ALS3 populations [8]. Expression of Tacr1 is far higher than that of Gpr83 in both ALS2 and ALS3 [8], and our findings in this study suggest that there are more Tacr1-expressing than Gpr83-expressing ALS cells in the LSN. It is therefore likely that cells belonging to one or more of these populations account for the difference in projections to the PBcl. The finding of a sizeable projection from both genotypes to the internal lateral parabrachial (PBil) nucleus by Choi et al [6] and in the present study presumably results from Tacr1 and Gpr83 expressing ALS neurons in the deep dorsal horn, including the lateral part of lamina V [21], since PBil receives input from spinal neurons located in the deep grey matter [36,37].

There were also significant differences between our findings and those of Choi et al [6], since we observed a similar pattern of labelling in the dorsal lateral parabrachial nucleus (PBdl) in the two genotypes, whereas Choi et al reported very sparse labelling in PBdl in the Tacr1^CreERT2^ mouse. Another important difference is that unlike Choi et al, we observed labelling in the thalamic VPL nucleus in both genotypes. This was expected, because VPL is known to be innervated by ALS neurons located in the superficial laminae [19]. The thalamic labelling reported by Choi et al resulted from an intersectional genetic strategy, in which the CreERT2 lines were crossed with a spinal cord selective Flp line (Lbx1^Flp^) and a Cre^ON^/Flp^ON^ reporter mouse, thus labelling axons of cells that expressed both recombinases. A possible explanation of the difference between our findings is therefore that the lamina I ALS cells did not contain sufficient levels of Flp in the Lbx1^Flp^ mouse.

## Conclusion

Our results indicate that among spinal projection neurons, expression of Tacr1 and Gpr83 is not restricted to ALS neurons in the SDH, but is found in ALS neurons in other locations as well as in projection neurons belonging to other tracts (PSDC and spinocerebellar pathways). They also show that even among ALS neurons in the SDH there is considerable overlap of expression of these two receptors. The finding of projections to VPL suggests that ALS cells expressing both of these receptors can contribute to the sensory-discriminative dimension of pain.

## List of abbreviations

ALS: anterolateral system
CTB: cholera toxin B subunit
CVLM: caudal ventrolateral medulla
IOM: medial nucleus of the inferior olive
LPAG: lateral part of periaqueductal grey matter
LPB: lateral parabrachial area
LSN: lateral spinal nucleus
MD: mediodorsal nucleus of thalamus
NK1r: neurokinin 1 receptor
NTS: nucleus of the solitary tract
PAG: periaqueductal grey matter
PoT: posterior triangular nucleus of the thalamus
PSDC: postsynaptic dorsal column pathway
SDH: superficial dorsal horn
VPL: ventral posterolateral nucleus of thalamus

## Acknowledgements

We are grateful to Mr Robert Kerr, Mr Iain Plenderleith and Ms Erin Dunn for excellent technical assistance, to Ms Aimi Razlan for help in some experiments, and to Dr D Ginty for the generous gift of Tacr1^CreERT2^ and Gpr83^CreERT2^ mice. The work was supported by the Wellcome Trust (grant number 219433/Z/19/Z) and the MRC (grant number MR/V033638/1).

## Authors’ contributions

WM, AMB and AJT designed the study. WM, ACD, EP, MY, RQ and MGM performed the experiments, and WM analysed the data. All authors contributed to the writing of the manuscript and approved the final version.

## Data availability statement

The datasets generated and analysed during the current study are available from the corresponding authors on reasonable request.

