## Supplementary material for "Expression of Tacr1 and Gpr83 by spinal projection neurons"

**Supplemental Material**

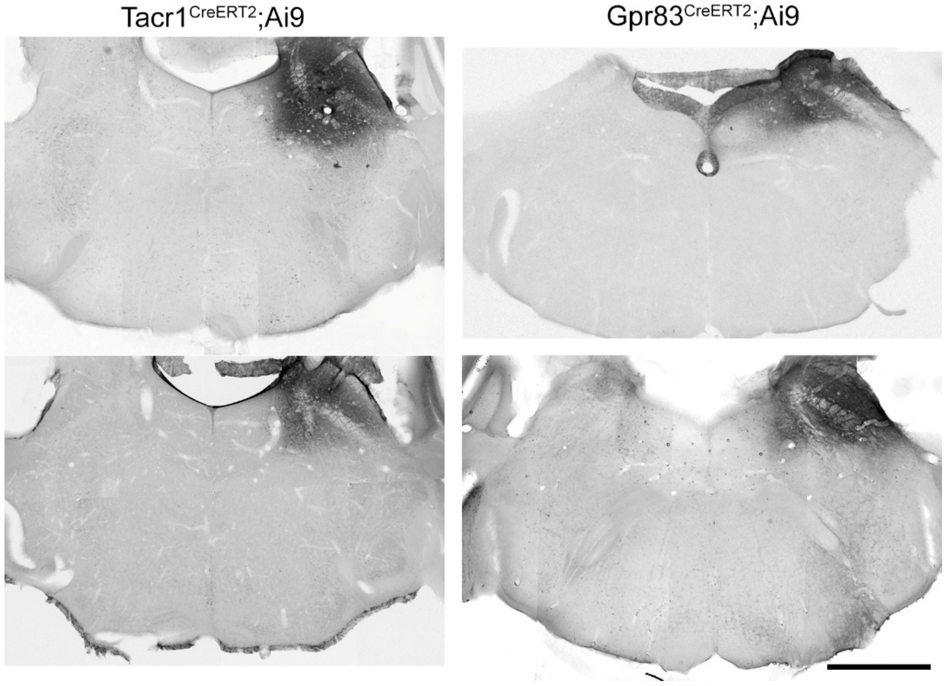


**Fig. S1**. Lateral parabrachial (LPB) injection sites in Tacr1$^{CreERT2}$;Ai9 and Gpr83$^{CreERT2}$;Ai9 mice. Cholera toxin B has been revealed with an immunoperoxidase method. Scale bar = 1 mm.

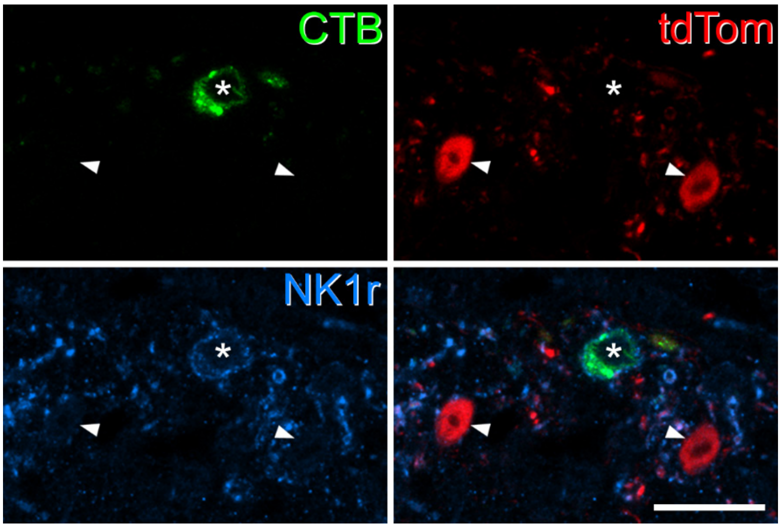


**Fig. S2**. Examples of the mismatch between tdTomato and NK1 receptor (NK1r) expression in the Tacr1$^{CreERT2}$;Ai9 mouse. Confocal images showing part of the superficial dorsal horn. Cholera toxin B (CTB) is shown in green, tdTomato (tdTom) in red, and NK1r immunoreactivity in blue. Arrowheads point to two CTB-negative cells with strong tdTomato signal that lack detectable NK1r. The asterisk indicates a neuron retrogradely labelled with CTB from the CVLM that is NK1r-immunoreactive, but does not contain tdTomato. Scale bar = 20 µm.

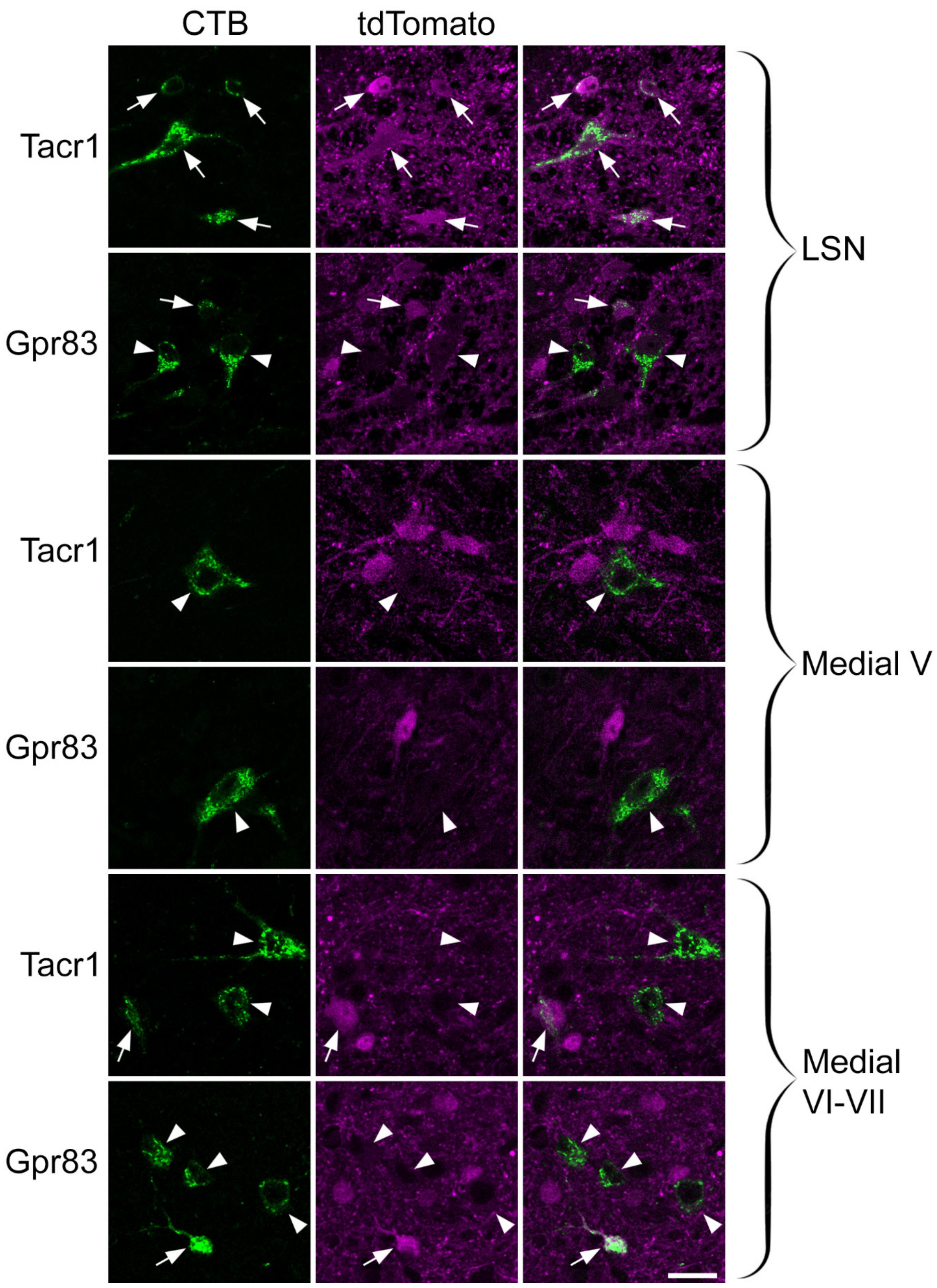


**Fig. S3.** TdTomato expression in retrogradely labelled neurons in regions other than the superficial dorsal horn. CTB (green) and tdTomato (magenta) are shown in Tacr1$^{CreERT2}$;Ai9 (Tacr1) and Gpr83$^{CreERT2}$;Ai9 (Gpr83) mice in the lateral spinal nucleus (LSN), the medial part of lamina V (Medial V) and the medial parts of laminae VI-VII (Medial V-VII). Arrows indicate CTB-labelled cells that contain tdTomato, and arrowheads those that lack tdTomato. Scale bar = 20 μm.

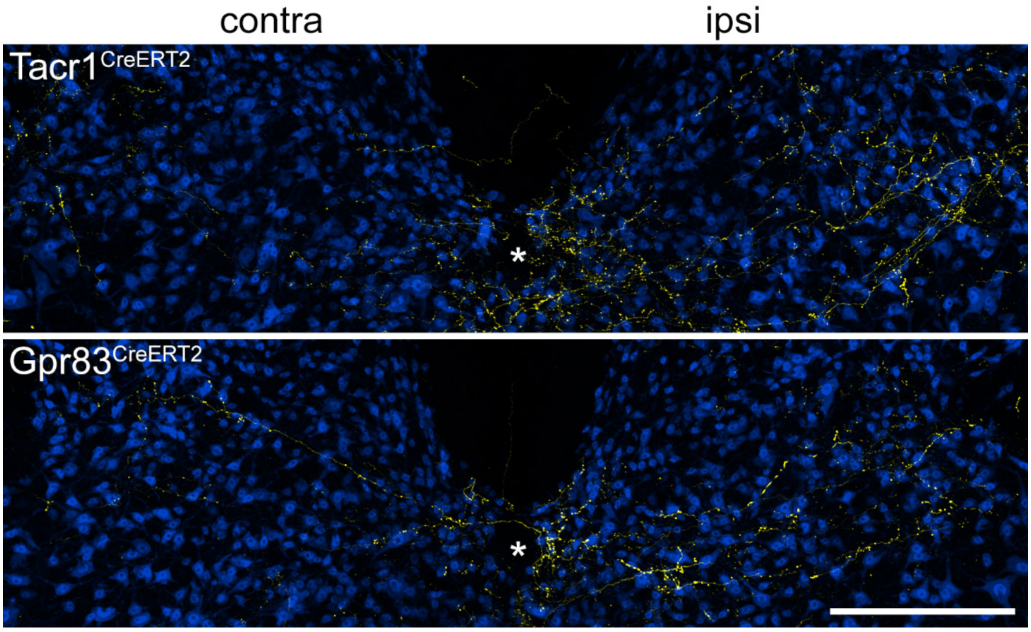


**Fig. S4.** TdTomato expression in axon collaterals in the spinal grey matter at the C7 level, following injection of AAV.Cre$^{ON}$.tdTomato into the L3, L4 and L5 segments of Tacr1$^{CreERT2}$ and Gpr83$^{CreERT2}$ mice. Sections have been immunoreacted to reveal tdTomato (yellow) and NeuN (blue). These images show higher magnification views of the regions outlined by boxes in Figure 3. Note the presence of numerous collateral branches, which are much more numerous on the side ipsilateral (ipsi) to the injection site, compared to the contralateral (contra) side. Scale bar = 20 μm.

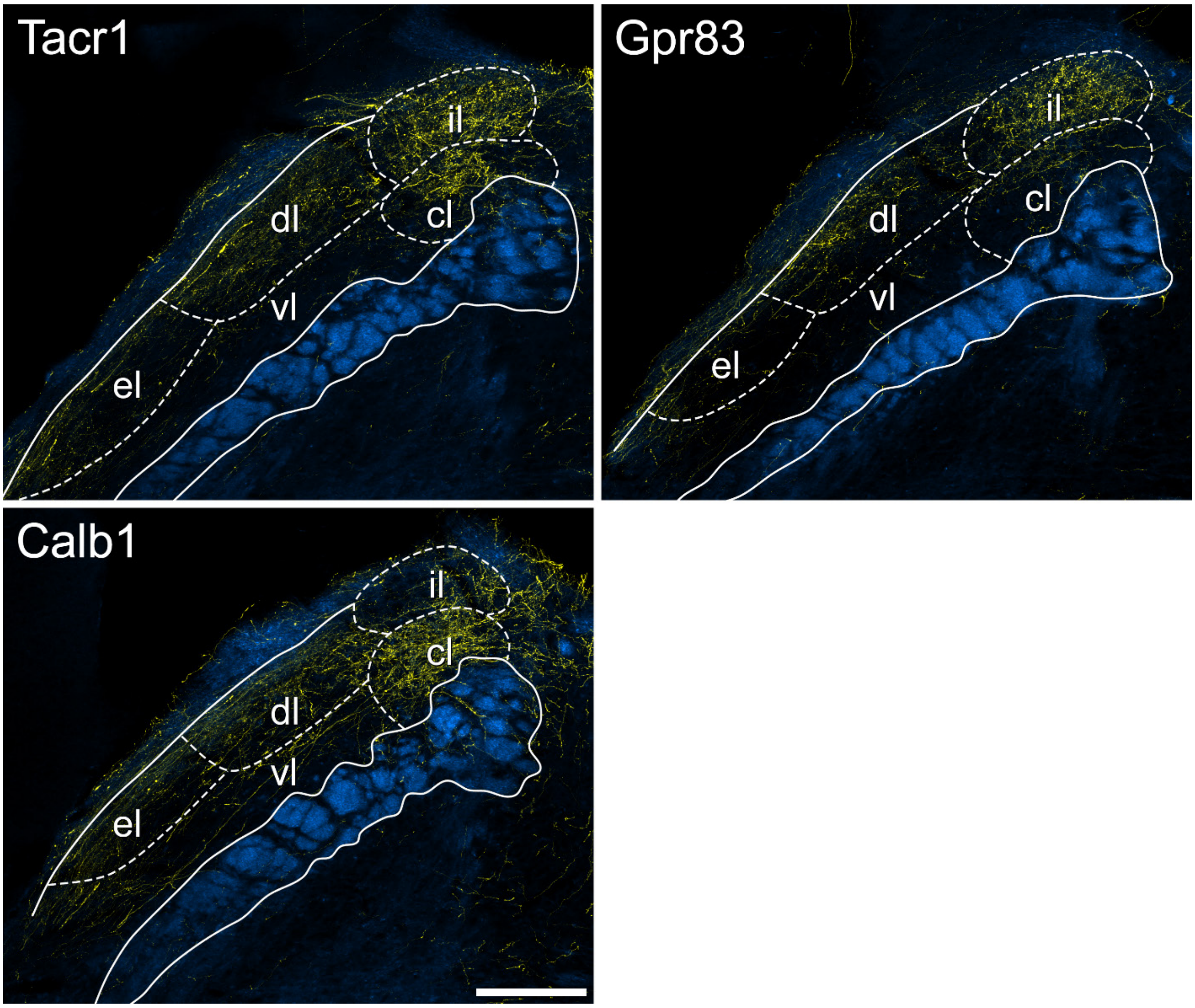


**Fig. S5.** TdTomato labelling in the lateral parabrachial (LPB) area in anterograde experiments that involved spinal injections of AAV.Cre$^{ON}$.tdTomato into a single segment (L3). Images show sections through the contralateral LPB scanned to reveal tdTomato (yellow), which is superimposed on a dark-field scan (blue) in Tacr1$^{CreERT2}$ (Tacr1), Gpr83$^{CreERT2}$ (Gpr83) and Calb1$^{Cre}$ (Calb1) mice. The pattern of axonal labelling in the Tacr1$^{CreERT2}$ and Gpr83$^{CreERT2}$ mice is very similar to that seen in mice of these genotypes that had received 3 spinal injections (shown in Fig 5), with labelling in el, dl and il in both genotypes. Axonal labelling in cl is far denser in the Tacr1$^{CreERT2}$ mouse compared to the Gpr83$^{CreERT2}$ animal. For comparison, we also illustrate anterograde labelling in a Calb1$^{Cre}$ mouse that had received a spinal injection of AAV.Cre$^{ON}$.tdTomato, and this has a dense plexus of tdTomato-positive axons in cl. Dense labelling in cl was also seen in 2 other Calb1$^{Cre}$ mice in which AAV.Cre$^{ON}$.tdTomato was injected into the L3, L4 and L5 segments. The superior cerebellar peduncle is outlined for each section, and approximate positions of LPB nuclei are illustrated: cl, central lateral; dl, dorsolateral; el, external lateral; il, internal lateral; vl ventral lateral. Scale bar = 200 μm.

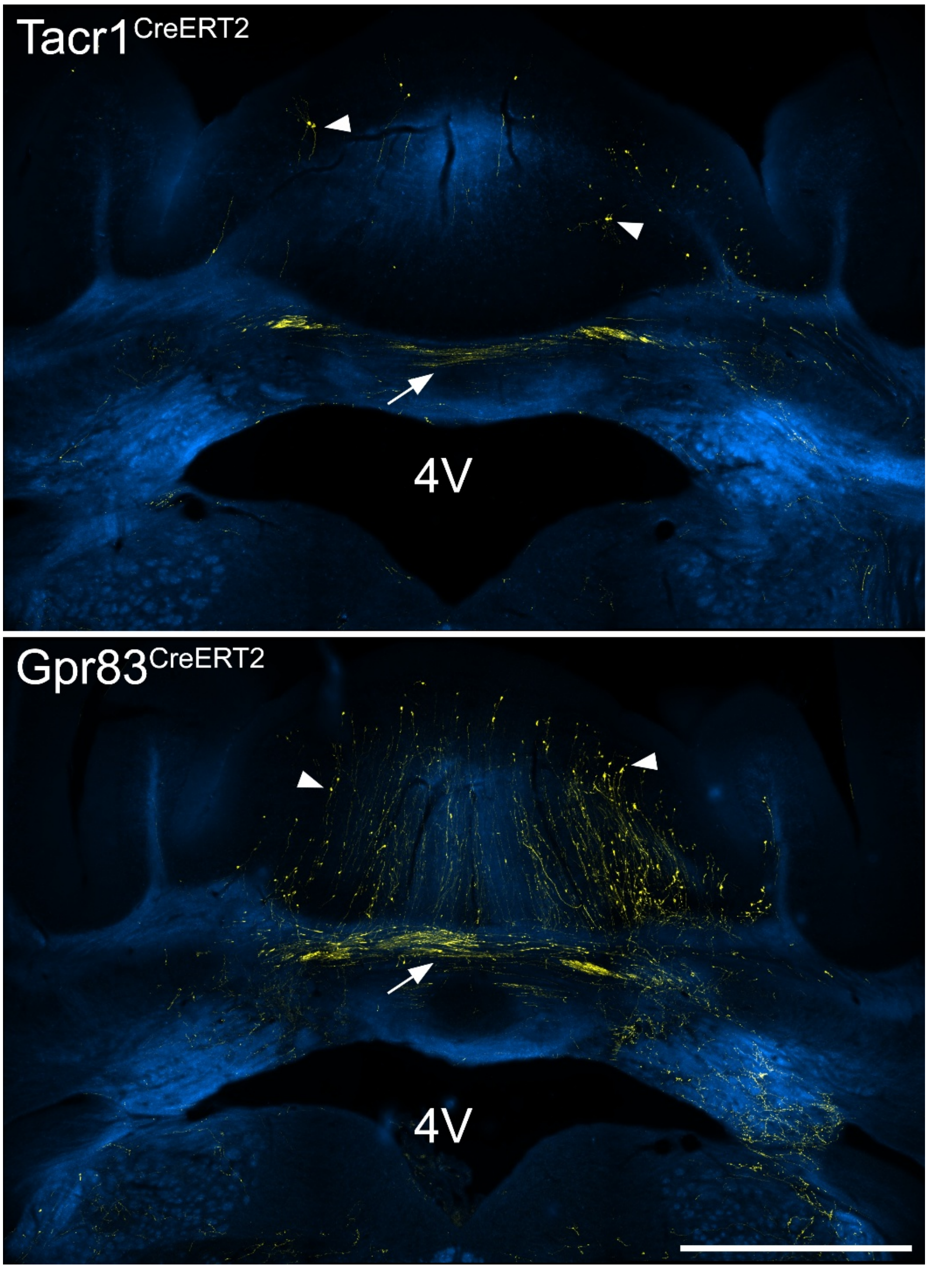


**Fig. S6.** TdTomato expression in axons in the cerebellum following injection of AAV.Cre$^{ON}$.tdTomato into the L3, L4 and L5 segments of Tacr1$^{CreERT2}$ and Gpr83$^{CreERT2}$ mice. Sections have been immunoreacted to reveal tdTomato (yellow), which is shown superimposed on a dark-field image (blue). In both cases, fibres cross the midline in the cerebellar white matter (cerebellar commissure, arrows). In addition, branches pass dorsally into the cerebellum, where they give rise to large boutons (some indicated with arrowheads), which presumably correspond to mossy fibre terminals. 4V, 4$^{th}$ ventricle. Scale bar = 1 mm.
